# Race specific differences in DNA damage repair dysregulation in breast cancer and association with outcome

**DOI:** 10.1101/2020.10.26.356006

**Authors:** Aloran Mazumder, Athena Jimenez, Rachel E Ellsworth, Stephen J Freedland, Sophia George, Matthew Bainbridge, Svasti Haricharan

## Abstract

**IMPORTANCE:** African American (AA) breast cancer patients have worse outcomes than Caucasian Americans (CAs). DNA damage repair (DDR) genes drive poor outcome in CA estrogen receptor (ER)+ breast cancer patients. Whether DDR genes similarly impact survival in AAs is unknown. Identifying AA-specific patterns of DDR dysregulation could change how we tailor predictive/prognostic biomarkers.

**OBJECTIVE:** To characterize DDR dysregulation in ER+ AA patient tumors and test associations with clinical outcome.

**DESIGN SETTINGS AND PARTICIPANTS:** Three independent tumor, and two normal breast datasets were analyzed. Tumor datasets: (1) GSE78958 (2) GSE18229 (3) The Cancer Genome Atlas (TCGA). Normal datasets: (4) GSE43973 (5) GSE50939.

**MAIN OUTCOME AND MEASURES:** Up/down-regulation of 104 DDR genes was assessed in AA samples vs CAs. Survival associations were assessed for genes dysregulated in multiple datasets.

**RESULTS:** Overall, RNA levels of single strand break repair (SSBR) genes were downregulated in AA tumors and double strand break repair (DSBR) genes were upregulated compared to CAs. While SSBR downregulation was mainly detected in tumors, DSBR upregulation was detectable in both tumor and normal breast AA samples. Seven specific DDR genes identified as dysregulated in AAs vs CAs in multiple datasets associated with poor survival. A subset of tumors with simultaneous dysregulation of homologous recombination and single strand break repair genes was enriched in AAs and had associated consistently with poor survival.

**CONCLUSION AND RELEVANCE:** Overall, these results constitute the first systematic analysis of differences in DDR regulation in AA ER+ tumors and normal tissue vs CAs. We identify a profile of DDR dysregulation enriched in AA patients, which associates with poor outcome. These results suggest a distinct molecular mechanism of DDR regulation in AAs that lays the groundwork for refining biomarker profiles by race and improving precision medicine for underserved populations.

## INTRODUCTION

African Americans (AAs) constitute a racial group with the highest mortality rate across cancer types. In AA women, breast, lung and colorectal are the three most common cancer diagnoses^1–4^. Breast cancer accounts for 32% of these diagnoses, making it one of the most predominant cancer types in AA women, as it is in Caucasian Americans (CAs)^4^. Although AA women typically have higher incidence of triple negative breast cancer than CAs, estrogen receptor positive (ER+) breast cancer remains the most commonly diagnosed subtype of breast cancer in AA women, as it is in CAs^2–4^. AA women with ER+ breast cancer present with higher tumor grade and more advanced disease that is more frequently luminal B (and therefore, resistant to standard endocrine therapies) than CAs. AA ER+ breast cancer patients are more likely to have poor clinical outcome with 42% higher mortality rate than CAs^5–9^.

There is consensus in the literature that environmental/socioeconomic factors contribute to poor breast cancer outcome in AAs^5,6,10,11^. These include reproductive factors and socioeconomic factors including access to healthcare^12–17^. However, even when these factors are controlled for, differences in presentation and outcome persist in AA ER+ breast cancer patients^15^. Therefore, it seems likely that ER+ tumor formation and progression in AA patients has a distinct molecular profile, as suggested by previous studies^18–20^. Understanding this profile is critical for developing precision medicine approaches tailored to this underserved population. Such efforts require comprehensive and race-specific molecular characterization of tumors.

AAs are, however, severely underrepresented in currently available datasets of patient tumors, which precludes comprehensive approaches to identify race-specific molecular drivers. This is true even of ER+ breast cancer, which is one of the most researched cancer types with large and multiple datasets comprised of whole genome/exome sequencing and gene expression from thousands of patient tumors. Although these datasets have too few AA tumors to power agnostic screens for drivers, it remains possible to use targeted approaches.

We, and others, have previously shown a link between DNA damage repair (DDR) dysregulation and endocrine therapy resistance in ER+ breast cancer^21–23^. Specifically, we identified causal links between defects in the MutL complex of mismatch repair (MMR), *CETN2* and *ERCC1* from nucleotide excision repair (NER) and *NEIL2* from base excision repair (BER) and resistance to endocrine therapies in ER+ breast cancer^23^. However, our studies generated molecular hypotheses from datasets where AA patient tumors were largely unrepresented (**Figure S1**). Of the datasets used in our original analysis, only one, TCGA, included data from AA patient tumors (n=43 AA vs n>800 CA)^23^. Therefore, our results reflect the DDR landscape of CAs. Since AAs have worse outcomes than CAs^2,12,24^, we hypothesize that resistance to endocrine therapy associated with higher frequency of dysregulation of DDR genes contributes to this outcome disparity. This is an especially relevant hypothesis since DDR pathways are molecular liaisons between external stressors (e.g. genotoxic stress, ageing) and regulation of common tumor phenotypes of growth, proliferation and apoptosis. Here, we test this hypothesis as described below.

## Material and Methods

### DDR gene set compilation

Gene sets for all DDR pathways analyzed (Mismatch repair, Nucleotide excision repair, Base excision repair, Non-homologous end joining, Fanconi Anemia and Homologous recombination) were derived from our previously published, curated DDR gene list (detailed in **Table S1**)^23^. Genes shared across different DDR pathways were not included.

### Datasets

Tumor datasets: Our first dataset (GEO78958, **Table S2**, referred to henceforth as dataset #1) had microarray data from 51 AA tumors with luminal (predominantly ER+) breast cancer, and 169 CA tumors^12^. Our second dataset (GSE18229, **Table S3**), referred to henceforth as dataset #2) had microarray gene expression data from 44 AA tumors and 85 CA tumors^25^. This dataset included tumors of all subtypes although ~70% of tumors were ER+/luminal. Our third dataset was the subset of ER+ tumors (irrespective of HER2 status) from TCGA and consisted of RNAseq gene expression and whole exome sequence data from 49 AA tumors and 449 CAs^26^ (**Table S4**). These restrictions were used to ensure that each dataset was predominantly ER+/luminal and that there was sufficient sample size of AA tumors. TCGA mutation data (downloaded March 2020) were obtained from cBioPortal. Gene expression from dataset #1 were available through the Gene Expression Omnibus (GEO, GSE78958). Gene expression from dataset #2^25^ were downloaded from dbGaP (downloaded May 2020) and for TCGA (downloaded March 2020) were obtained from cBioPortal. TCGA survival outcomes were downloaded from cBioPortal^27^ (downloaded May 2020). Standard cutoffs of mean+/-1.5× SD were used on the RNA data to identify “High” and “Low” subsets respectively in each dataset.

Normal datasets: Our normal breast tissue dataset (GSE43973)^28^, had microarray data from 12 AA women and 98 CA women. Our tumor adjacent normal tissue dataset (GSE50939)^29^ had microarray data from 14 AA patients and 52 CA patients.

### Enrichment analysis

For RNA analysis, p-values were obtained by comparing each gene between AA and CA tumors using Wilcoxon Rank Sum test. These p-values were rank-ordered and q-values were computed. All genes with q<0.25 (p<0.1) were considered candidates in each dataset. Candidate genes with RNA dysregulation in the same direction across two datasets were curated to form the final list of genes for downstream analyses. For overall patterns of RNA dysregulation in the union of the three datasets, each gene was included only once if it was dysregulated in the same direction in multiple datasets but included twice if it was dysregulated in opposite directions in multiple datasets. Fisher’s exact test determined p-values for overall patterns of up and down-regulation.

For mutation analysis, any gene with 2% incidence (i.e. ≥1 mutation in AA tumors) was considered. All non-synonymous mutations were included irrespective of category (i.e. missense, nonsense, frameshift, etc) or predicted pathogenicity. Expected rates of mutation frequency were calculated based on total number of mutations identified in the entire patient population and compared to observed rates in AA and CA tumors respectively. Fisher’s exact test determined p-values by comparing observed to expected frequency of each mutation.

For functional pathway analysis, functional category was determined based on literature searches for each gene of each pathway used in this targeted analysis (**Table S1**). Each gene was then assigned to sensor, scaffold, DNA repair and downstream effector categories based on a literature search. If a gene fell into two functional categories, it was considered in statistical analyses of each category in turn. The number of candidate genes in each category was compared to the total number of genes in that category using a Fisher’s exact test.

For proliferation analyses in **Figure 3**, *MKI67* (gene name for Ki67) RNA levels were used and the top 20% and bottom 20% of *MKI67* expressing tumors were considered as “High” and “Low” proliferators respectively. Fisher’s exact test determined p-values.

### Survival analysis

For univariate and multivariate analyses, all tumors with associated survival data in each dataset were used, with restriction to luminal A/B tumors in dataset #1 and ER+ tumors in TCGA. Outcome measures used were disease-free survival for dataset #1 and TCGA, and recurrence-free survival in dataset #2 in **Figure 1**, and in **Figures 4** and **S5**, disease-specific survival for dataset #1, recurrence-free survival for dataset #2 and overall survival for TCGA. These different outcome measures were used because they had the largest sample size associated with them in each dataset. Factors included in multivariate analyses were PAM50 status and tumor stage for dataset #1, PAM50, tumor size and nodal status for dataset #2 and PR/HER2 status and tumor stage for TCGA. For all datasets, race was included as a categorical, and age as a continuous, variable. Only samples with survival metadata were included in the analysis.

**Figure 1:**
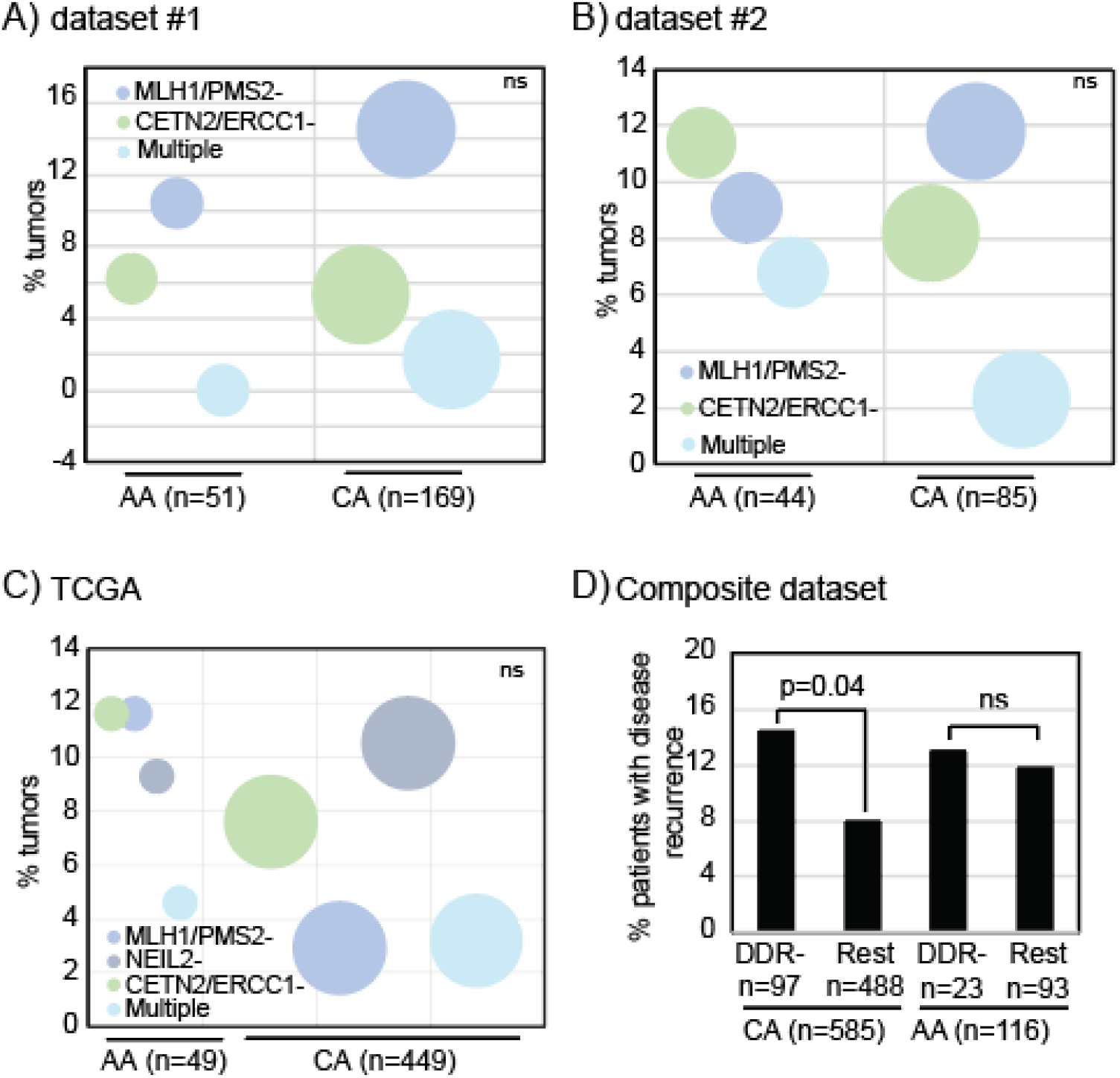
Downregulation of DDR genes that associate with poor outcome in CA ER+ breast cancer patients do not associate with poor outcome in AAs. (A-C) Bubbleplots representing % tumors from AA or CA patients with low RNA levels (mean-1.5*std devn) of DDR genes belonging to nucleotide excision repair (NER: *CETN2*, *ERCC1*), mismatch repair (MMR: *MLH1, PMS2*), or both (Multiple) in dataset #1 (A), dataset #2 (B) and TCGA (C). (D) Column graph depicting cumulative % of AA and CA patients with disease recurrence, from all three datasets, whose tumors either had detectable dysregulation of any DDR genes or did not. Pearson’s Chi Squared test determined all p-values. Associated descriptive characteristics of each of the three datasets in **Tables S2-4**.

### Statistical analysis

Missing data were imputed with “NA” from mutation, expression, and survival data analysis. Samples classifying for more than one category (e.g. SSBR and DSBR dysregulation) were treated as separate set for statistical comparisons. Two-tailed Wilcoxon rank sum tests were used for two-sample tests of comparing continuous data and Pearson’s Chi Square test (or Fisher’s Exact test) was used for comparing categorical data. Log rank test calculated p-values for survival analyses and Cox regression determined proportional hazards.

## Results

### Defects in previously identified DNA repair genes do not associate with outcome in African American patients

We first investigated the frequency with which DDR genes known to induce endocrine therapy resistance when lost (MMR: *MLH1* and *PMS2*, NER: *CETN2* and *ERCC1*, and BER:*NEIL2*)^23^ are downregulated in AA tumors from three selected datasets **(Figure S2).** In all three datasets (**Table S2-4**), frequency of downregulation of RNA levels of these DDR genes was similar between AA and CA tumors with no demonstrable enrichment by race **(Figures 1A-C).** We next tested whether downregulation of these DDR genes associated with poor disease-free survival in AA patients, as it did in CAs^23^. Because of the small sample size of AA patients in each dataset, we combined data from all three datasets for this analysis. There was no significant difference in disease recurrence between AA patients whose tumors had downregulation of these DDR genes vs patients whose tumors did not downregulate these genes (14% vs 12% disease recurrence, **Figure 1D).** In contrast, and as expected, we observed a statistically significant increase in disease recurrence in CAs with 14% of patients whose tumors downregulated these DDR genes having disease recurrence compared to only 8% of CAs (p=0.04) whose tumors did not detectably downregulate these DDR genes **(Figure 1D).** Overall, this analysis identified no enrichment in AA ER+ tumors for downregulation of the four DDR genes previously identified as driving poor outcome in CAs.

### Landscape of DNA repair dysregulation in African American tumors

To identify whether AA tumors have a distinct pattern of DDR dysregulation, we compared RNA levels of 104 DDR genes from six principle DDR pathways (**Table S1**) between AA and CA tumors in each of three independent datasets: dataset #1 (**Table S2**), dataset #2 (**Table S3**) and TCGA (**Table S4**). From the union of the three datasets, 67 genes were either up- or downregulated in AA tumors relative to CAs (q<0.25, **Figure S3).** SSBR genes were enriched in this gene set with 80% of NER genes and 90% of BER genes being either up- or down-regulated in AA tumors relative to CAs in at least one dataset (**Figure 2A**). Fanconi Anemia (FA) genes were similarly heavily represented (90% of genes making the cut-off). The remaining pathways had 50-60% of their genes represented **(Figure 2A)**. Overall, SSBR genes (NER, BER and MMR) had lower RNA levels in AA tumors relative to CAs, while DSBR genes (FA and HR) had higher RNA levels (**Figure 2B**). Non-homologous end joining (NHEJ) was the only DSBR pathway where gene expression was preferentially downregulated in AA tumors **(Figure 2B).** We next identified the list of genes dysregulated in the same direction in at least two of the three datasets analyzed. This gene list consisted of three BER genes *(NEIL3, XRCC1, PARP1*), one NER gene (*MNAT1*), one NHEJ gene (*XRCC4*), one FA gene (*FANCE*) and two HR genes (*NBN and BRCA1*) **(Figure 2C)**. Consistent with overall patterns of dysregulation, FA and HR (DSBR) genes were upregulated, *XRCC4* (NHEJ) was downregulated, and two of the three BER genes were downregulated **(Figure2C**, **FigureS4, FigureS5)**.

**Figure2:**
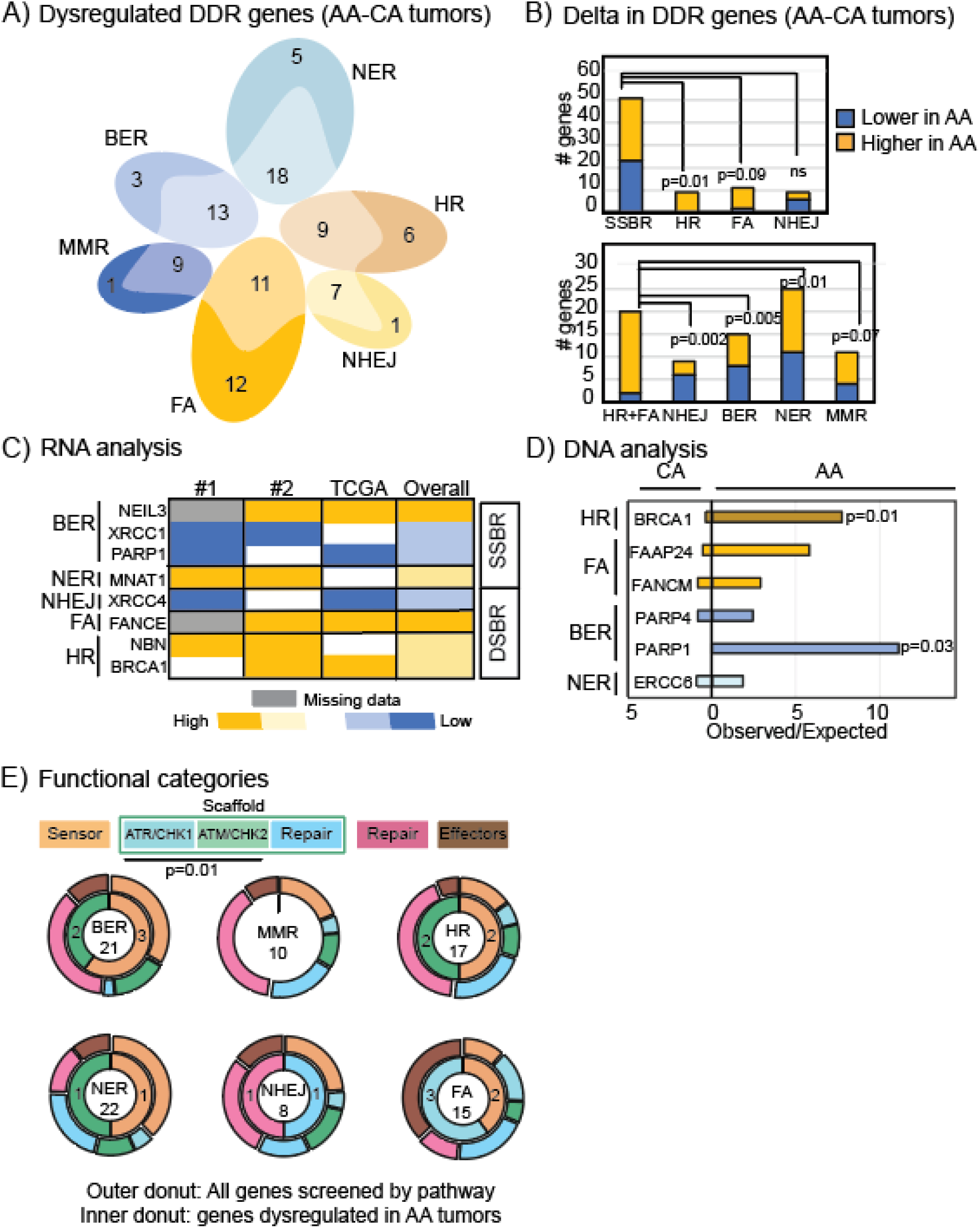
AA tumors have a distinct pattern of DDR dysregulation. (A) Venn diagram showing proportion of genes from each of six DDR pathways (listed in **Table S1**) that are significantly (q<0.25) dysregulated in AA tumors relative to CAs in at least one of three datasets analyzed. Schematic of analysis in **Figure S2** Associated **Figure S3** presents full list of DDR genes. (B) Stacked column graphs representing number of DDR genes that are up (yellow) or down (blue)-regulated by pathway. Pearson’s chi-squared test determined p-values. (C) Heatmap showing candidate genes that are either up-(yellow) or down-regulated (blue) in at least two datasets as assessed by q-value analysis based on p-values derived from Wilcoxon Rank Sum tests. Associated **Figure S4-S5** presents boxplots of each candidate gene described in the heatmap. (D) Bar graph showing observed/expected ratio for mutational frequency of all DDR genes mutated at least once in AA patient tumors from TCGA. Observed/expected ratio for CA tumors are presented by bars to the left of the median line, while the ratio for AAs is presented to the right. Pearson’s chi-squared test determined p-values. Associated **Figure S6** presents additional data on DDR gene mutations in AA and CA tumors. (E) Layered donut plot depicting proportional representation of functional DNA repair categories in the 104 DDR genes screened (outer donut) and in the list of 12 genes identified in (C) and (D) (inner donut). Fisher’s exact test determined p-value. ATM/Chk2 and ATR/Chk1 scaffolds were combined to represent cell cycle checkpoint scaffolds, as a distinct category from repair-scaffolds. Full list of genes in functional categories presented in **Table S1**. NER, Nucleotide excision repair, BER, Base excision repair, MMR, mismatch repair, NHEJ, non-homologous end joining, FA, Fanconi anemia, HR, homologous recombination, SSBR, single strand break repair, DSBR, double strand break repair. #1, dataset #1, #2, dataset #2.

Using whole exome sequence data from TCGA, we next tested whether any DDR genes were enriched for mutations in AA tumors. Other than *PARP1*, none of the DDR genes differentially regulated in AA tumors from the RNA analysis were mutated in AA tumors (**Figure S6A**). Overall, 16% of AA tumors had mutations in at least one DDR gene, compared to only 3% of CA tumors (p<0.001), with specific enrichment for mutations in genes from BER and HR pathways (**Figure S6B**). In total, six genes: *ERCC6* (NER), *PARP1* and *PARP4* (BER), *FANCM* and *FAAP24* (FA) and *BRCA1* (HR) had increased mutational frequency in AA tumors vs CAs, of which, *PARP1* and *BRCA1* (also identified in RNA analysis) reached statistical significance (p=0.01, p=0.03) respectively **(Figure 2D).** Mutations in any DDR gene associated with significantly worse disease-free survival in AA patients (HR=4.12, p=0.02, **Figure S6C**).

To test whether any functional patterns emerged in the DDR genes identified as differentially regulated at RNA level/mutated in AA tumors, we assessed distribution of these genes in the cascade of events involved in each DDR pathway. We categorized genes based on their primary function as Sensor, Scaffold, Repair or Effectors (**Table S1**). Sensors sense the presence of specific types of DNA damage, Scaffold proteins serve to stabilize and activate other proteins at the site of damaged DNA, while Repair and Effector proteins are directly involved in repairing damaged DNA. Scaffold proteins were further categorized as cell cycle checkpoint kinase scaffolds that stabilize and activate ATM/Chk2 and ATR/Chk1 kinases or repair scaffolds that stabilize and activate downstream DNA repair proteins. Components of any DDR pathway predominantly function as sensor or repair/effector proteins, with only 15-20% of the pathway proteins identified as cell cycle checkpoint scaffolds (**Figure 2E**). However, the majority of DDR genes dysregulated in AA tumors are ATM/ATR scaffolds. This is in keeping with previous findings that DDR genes contribute to endocrine therapy resistance by dysregulating cell cycle checkpoint activation in response to endocrine therapy ^22,23,30(p4)^

### Persistent upregulation of double strand break repair genes in normal breast tissue from African American women

To understand whether differences in DDR dysregulation patterns in AA tumors are driven by endogenous differences in gene expression of breast tissue, we used microarray gene expression data from two publicly available datasets: GSE43973 with samples from reduction mammoplasty (12 AA and 98 CA women) and GSE50939 with samples from tumor-adjacent normal tissue (14 AA and 67 CA patients)^28,29^. Overall, the predominant difference in DDR gene expression between AA and CA breast tissue was upregulation of DSBR genes, while SSBR genes were rarely different between these two cohorts **(Figure 3A-D).** Even NHEJ genes which are preferentially downregulated in AA tumors were upregulated in the normal breast (**Figure 3B)**. The number of dysregulated genes in the normal breast of AAs vs CAs was far fewer than that seen in tumors (**Figure 3B-D**).

**Figure3:**
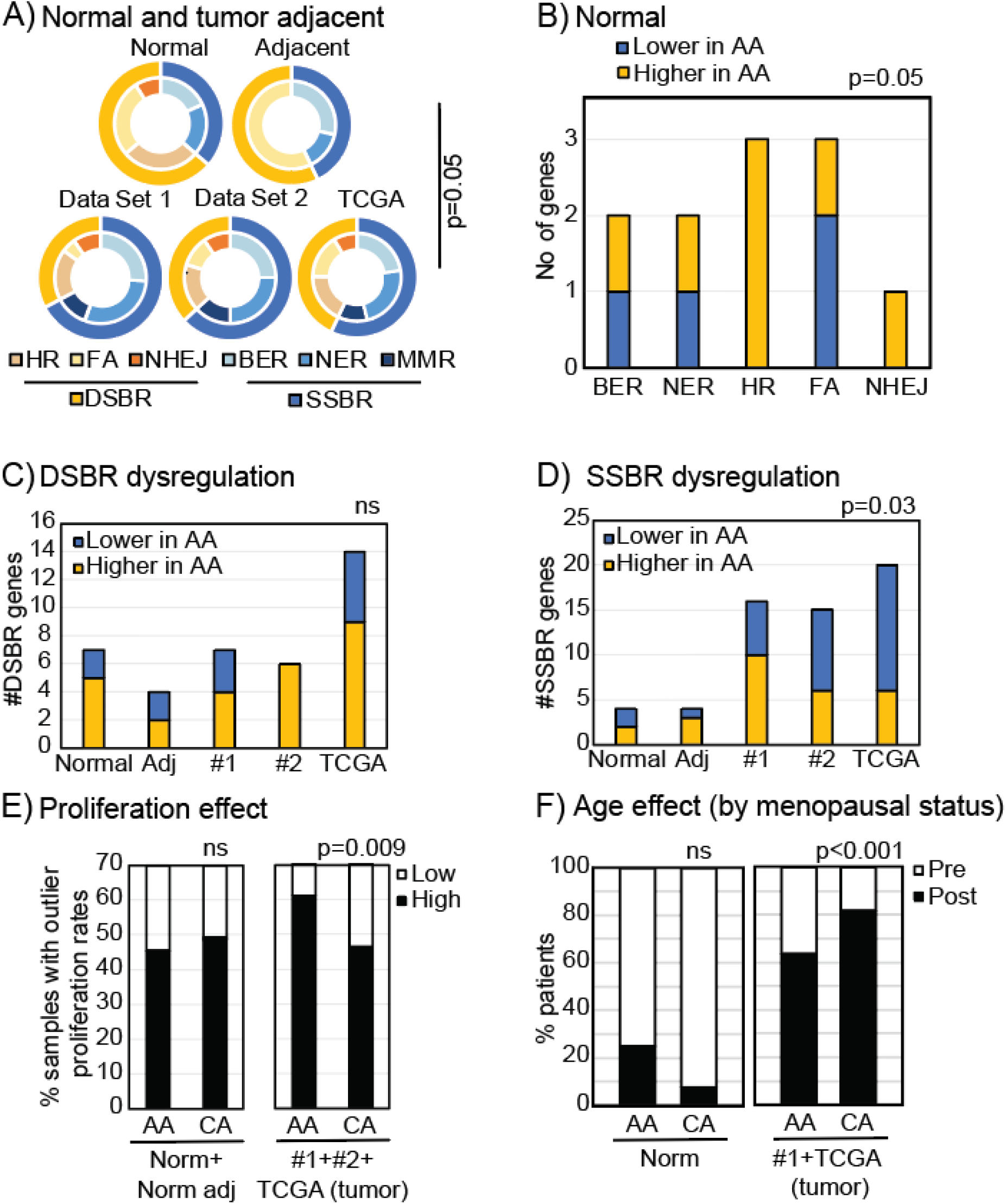
Differences in the DDR landscape of AA vs CA normal breast tissue. (A). Nested donut plots representing proportion of SSBR (blue) vs DSBR (yellow) genes (outer donut) and proportion of genes from each pathway within SSBR and DSBR categories (inner donut) that were significantly (p<0.1) up- or down-regulated in AA vs CA normal breast samples. Wilcoxon Rank Sum test determined p-values. Distribution of DDR dysregulation in tumor datasets analyzed in **Figure 2** is presented for comparison. (B) Stacked column graph representing # genes by pathway that are significantly (p<0.1) up (yellow) or down (blue) regulated in AA relative to CA normal breast tissue. (C-D) Stacked column graphs summarizing numbers of DSBR (C) and SSBR (D) genes that are either up- (yellow) or down-(blue) regulated in normal (Normal and Adj) vs tumor (#1, #2, TCGA) datasets. (E-F) Stacked column graphs representing proportion of high and low proliferating samples (E) and pre- and post-menopausal (F) AA and CA women in normal vs tumor datasets. High proliferating samples reflect the top 20% of tumors ranked by gene expression of *MKI67*, a marker of proliferation from high to low, while low proliferating samples reflect the bottom 20th percentile. Pearson’s chi-squared test determined all categorical p-values. NER, Nucleotide excision repair, BER, Base excision repair, MMR, mismatch repair, NHEJ, non-homologous end joining, FA, Fanconi anemia, HR, homologous recombination, SSBR, single strand break repair, DSBR, double strand break repair. #1, dataset 1, #2, dataset 2, Adj, tumor-adjacent normal, Norm, normal.

We next tested whether baseline upregulation of DSBR genes in AA normal breast tissue and preferential SSBR downregulation in tumors associates with higher proliferation in AA tumors or younger age of AA women in our datasets. Both proliferation and age are known to affect DDR gene expression and AA breast cancer patients present with more proliferative tumors at a younger age^19^. In combined data from normal and tumor adjacent normal datasets, using gene expression of *MKI67* as a proliferative index, we found no differences between AA and CA women (45% vs 49% were highly proliferative, **Figure 3E**). However, in the combined tumor datasets, proliferation index was significantly higher in AA tumors (61% of tumors) compared to CAs (46%) p=0.009 **(Figure 3E**). Therefore, it is possible that the loss of SSBR gene expression in AA tumors is associated with higher proliferation. A similar trend was detected when age was considered using menopausal status as a surrogate. While we found no statistically significant difference in age in women represented in normal datasets, the number of post-menopausal tumor samples was significantly higher in CA patients (81%) compared to AAs (63%), p<0.001 **(Figure 3F)**. Neither of these parameters explains the higher DSBR gene expression observed in normal AA breast tissue.

### African American-specific DNA repair dysregulation associates with poor patient outcomes

To assess prevalence of DDR dysregulation in AA patients, we tested the proportion of tumors with dysregulation of any of our DDR genes (DSBR: HR – upregulation of *NBN, BRCA1*, FA – upregulation of *FANCE*, NHEJ – downregulation of *XRCC4*; SSBR: BER – downregulation of *PARP1*, *XRCC1*, upregulation of *NEIL3*, NER – upregulation of *MNAT1*) in AAs vs CAs. In datasets #1 and #2, AA tumors dysregulated at least one SSBR gene with a >2-fold increase in frequency relative to CAs (p=0.01 in each case, **Figure S7A-B**). In dataset #2 alone, AA tumors also dysregulated at least one DSBR gene with significantly increased frequency relative to CAs (**Figure S7B**). In TCGA, frequency of SSBR and DSBR gene dysregulation was comparable between AA and CA tumors (**Figure S7C**). However, in all three datasets, we observed a subset of tumors that were significantly enriched in AA patients. This subset simultaneously dysregulated SSBR and DSBR genes, and accounted for >10% of AA tumors, but <3% of CA tumors, >3-fold enrichment (**Figure S7A-C**). This subset also associated with significantly worse outcome in datasets #1, #2 and TCGA, irrespective of whether patients were AA or CA (**Figure S7D-F**).

To further understand this subset, we parsed individual gene dysregulation. In all three datasets, we observed simultaneous dysregulation of one of the two HR genes, *NBN* or *BRCA1*, with different SSBR genes (an HR/SSBR subset). The NER pathway gene, *MNAT1* was implicated in this simultaneous dysregulation in all three datasets (**Figure 4A-C**). In datasets #1 and #2, the HR/SSBR subset was significantly enriched in AAs (6-8%) relative to CAs (1-3%) (**Figure 4A-B**), while in TCGA, an FA/SSBR subset was significantly enriched (10% vs 2%, p<0.001), **Figure 4C**). This is likely because of all three datasets, TCGA had the best representation of FA genes (which were largely missing from datasets #1 and #2), and the most frequent upregulation of *FANCE* in AA tumors.

**Figure4:**
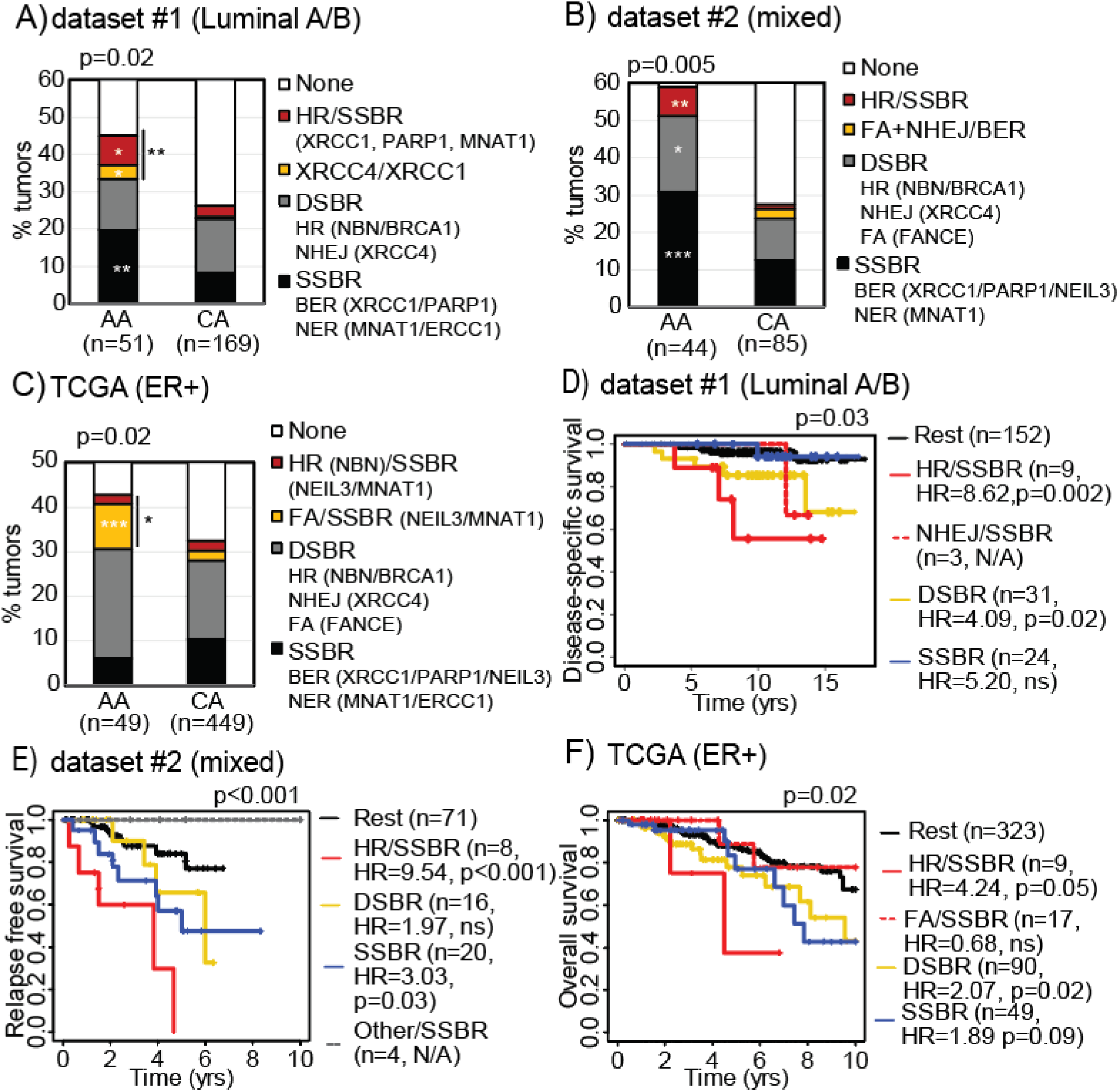
DDR genes enriched for dysregulation in AA tumors associate with worse survival. (A-C) Stacked columns representing % of AA vs CA tumors with dysregulation of any one of the DDR genes from the specified pathways in dataset #1 (A), dataset #2 (B) and TCGA (C). Pearson’s chi-squared test determined p-values. Associated data presented in **Figure S7A-C**. (D-F) Kaplan-Meier curves representing disease-specific survival in dataset #1 (D), relapse-free survival in dataset #2 (E) and overall survival in TCGA (F) associated with patients whose tumors had dysregulation of specified DDR genes by pathway relative to tumors that did not. Log rank test determined p-values. Associated data in **Figure S7D-F**, and Cox Regression Proportional Hazards assessment in **Tables S5-7**. HR in survival curves, hazard ratio, p<0.05, *; p<0.01, **; p<0.001, ***.

Since the HR/SSBR subset was represented in all datasets analyzed and was significantly enriched in AA tumors in two of three datasets, we next tested its association with outcomes. We broke each dataset up into patients whose tumors dysregulated any SSBR gene (downregulation of *XRCC1/XRCC4/PARP1*, upregulation of *NEIL3/MNAT1*) alone and any DSBR gene (upregulation of *NBN/BRCA1/FANCE*) alone. Tumors with simultaneous dysregulation of SSBR and DSBR genes were divided into those with simultaneous upregulation of either HR gene, *NBN/BRCA1*, with dysregulation of any SSBR gene (downregulation of *XRCC1/PARP1*, upregulation of *NEIL3/MNAT1*). We compared associations with survival outcome of each of these groups relative to patients whose tumors had no detectable dysregulation of any of these genes. The only patient group that consistently and significantly associated with worse disease-specific survival (dataset #1, HR=8.62, p=0.002, **Figure 4D**), relapse-free survival (dataset #2, HR=9.54, p<0.001, **Figure 4E**) and overall survival (TCGA, HR=4.24, p=0.05, **Figure 4F**) was the HR/SSBR subset. On the other hand, the simultaneous dysregulation of non-HR DSBR candidates with any SSBR gene did not associate with survival outcomes in any dataset (**Figures 4D-F**).

Next, we conducted Cox Proportional Hazards assessment to understand the effect of confounding factors (tumor stage or tumor grade/nodal status, PAM50 or PR/HER2 status, age and race) on associations between DDR dysregulation and survival. In datasets #1 and #2, but not in TCGA, we found that association of HR/SSBR tumors with poor survival remained significant even in a proportional hazards assessment (**Tables S5-7**). Age and race did not remain significant in any of the three datasets, suggesting that the unique candidate gene set identified by analyzing AA patient tumors is likely a poor prognostic factor regardless of race, although significantly enriched in AAs.

## Discussion

DNA repair proteins are natural molecular conduits between external stimuli and cellular response. Exposure to genotoxins or hypoxia, for instance, can induce a cell to up- or down-regulate its DDR signaling^31^. Not only do DDR proteins repair damaged DNA they also activate cell cycle checkpoints and engage apoptotic pathways^32^. Therefore, DDR pathways make an attractive starting point for understanding how molecular factors that translate environmental stimuli into cellular phenotypes may be altered by race/ethnicity.

However, it is important to outline several caveats of this study. Firstly, each of the tumor datasets analyzed in this study had ≤51 AA patient tumors. This low sample size likely resulted in high false negative rates. Secondly, the three datasets used had different, although overlapping, composition of patient tumors. These differences undoubtedly limited our ability to interpret nuances of the association of DNA repair dysregulation with survival. For instance, upregulation of any DSBR gene associated with poor patient outcome in dataset #1 (HR=4.09, p=0.02, **Figure 4D**) and in TCGA (HR=2,07, p=0.02, **Figure 4F**), both of which were restricted to ER+ tumors, but not in unrestricted dataset #2. In contrast, dataset #2 was the only dataset where patients whose tumors had SSBR dysregulation alone had significantly worse relapse-free survival (HR=3.03, p=0.03, **Figure 4E**). Whether this is because SSBR dysregulation has a particularly strong association with relapse-free survival, or if it is because this dataset included 30% of ER-tumors, is uncertain, but an interesting question for further study.

To some extent these challenges are circumvented by Cox Proportional Hazards analysis. But discrepancies in the results of the Cox analysis emphasize the issues caused by dataset heterogeneity. In TCGA, for example, which had the highest representation of FA genes amongst the three datasets, simultaneous HR/SSBR dysregulation was not an independent prognostic factor in Cox analysis (**Table S7**). In contrast, DSBR dysregulation and HER2 status remained significant prognostic factors in TCGA after proportional hazards assessment (**Table S7**), perhaps indicating a confounding role for HER2 positivity in the association between DNA repair and overall survival. Again, tumor stage remained a significant prognostic factor for dataset #1, our most homogeneous dataset (**Table S5**), while SSBR dysregulation and PAM50 status remained significant in dataset #2, our most heterogeneous dataset (**Table S6**).

Normal datasets had even lower AA sample size, thereby enabling pathway-level analysis but not gene resolution. Finally, we only had access to a single dataset, TCGA, for mutation data. Although we identified significant enrichment for mutations in two genes previously implicated in race-based molecular differences, *PARP1* and *BRCA1* ^2,33^ we suspect that other genes would attain significance given sufficient sample size. Overall, a more thorough investigation of the nuances of DNA repair dysregulation in AA tumors is warranted in larger discovery datasets that are homogeneous and sufficiently powered to resolve these discrepancies.

In spite of these caveats, several novel and interesting molecular discoveries were made through these analyses, which we hope will spur efforts to generate larger, better annotated AA datasets. This study provides the first evidence of a spectrum of DDR dysregulation in ER+/HER2− breast tumors enriched in AA patients and associating with poor outcome. Of note, no MMR gene was consistently downregulated in AA tumors across our three datasets. Given our previous discovery of *MLH1*/*PMS2* downregulation as a key mediator of poor outcome in CA ER+ breast cancer patients^22^, it is possible that this molecular driver is shared between AA and CA patients, and is therefore not significantly further dysregulated in AAs. Most strikingly, all three datasets interrogated in this study demonstrated significant enrichment for simultaneous SSBR/DSBR dysregulation in AA tumors relative to CAs. These data highlight the existence of an AA-specific DDR dysregulation pattern in ER+ tumors that is likely prognostic and may be predictive of endocrine therapy response. Further experimental investigation is warranted. Overall, this study presents a roadmap for developing AA-specific biomarkers to prevent poor outcomes. This study could also have implications for other endocrine-related cancer types where AA-specific outcome disparities are observed^34–38^.

## Supporting information

Supplementary Mazumder_et_al_2020

## Acknowledgements

We acknowledge Drs. Melissa Troester and Meenakshi Anurag for generously sharing resources for the completion of this work. Work in this study was funded by Department of Defense Breast award (W81XWH-18-1-0034 to SH), NCI K22 Career Development award (CA229613 to SH), and Susan G. Komen Career Catalyst (CCR18548157 to SH). Authors declare no conflicts of interest.

## References

1. Furberg H, Millikan R, Dressler L, Newman B, Geradts J. Tumor characteristics in African American and white women. Breast Cancer Res Treat. 2001;68(1):33–43. doi:10.1023/A:1017994726207

2. Gao R, Price DK, Sissung T, Reed E, Figg WD. Ethnic Disparities in Americans of European descent versus Americans of African descent related to Polymorphic ERCC1, ERCC2, XRCC1 and PARP1. Mol Cancer Ther. 2008;7(5):1246–1250. doi:10.1158/1535-7163.MCT-07-2206

3. Haddad SA, Lunetta KL, Ruiz-Narváez EA, et al. Hormone-related pathways and risk of breast cancer subtypes in African American women. Breast Cancer Res Treat. 2015;154(1):145–154. doi:10.1007/s10549-015-3594-x

4. Breast Cancer Rates Among Black Women and White Women | CDC. Published January 31, 2019. Accessed October 20, 2020. https://www.cdc.gov/cancer/dcpc/research/articles/breast_cancer_rates_women.htm

5. Rauscher GH, Silva A, Pauls H, Frasor J, Bonini MG, Hoskins K. Racial disparity in survival from estrogen and progesterone receptor-positive breast cancer: implications for reducing breast cancer mortality disparities. Breast Cancer Res Treat. 2017;163(2):321–330. doi:10.1007/s10549-017-4166-z

6. Menashe I, Anderson WF, Jatoi I, Rosenberg PS. Underlying Causes of the Black–White Racial Disparity in Breast Cancer Mortality: A Population-Based Analysis. JNCI J Natl Cancer Inst. 2009;101(14):993–1000. doi:10.1093/jnci/djp176

7. Ma H, Lu Y, Malone KE, et al. Mortality risk of black women and white women with invasive breast cancer by hormone receptors, HER2, and p53 status. BMC Cancer. 2013;13:225. doi:10.1186/1471-2407-13-225

8. Ren Y, Black DM, Mittendorf EA, et al. Crossover Effects of Estrogen Receptor Status on Breast Cancer-Specific Hazard Rates by Age and Race. PLOS ONE. 2014;9(10):e110281. doi:10.1371/journal.pone.0110281

9. Menendez-Tuckman AT, Raventos-Suarez C. Re: Tumor biologic factors and breast cancer prognosis among white, Hispanic, and black women in the United States. J Natl Cancer Inst. 1994;86(17):1352–1353. doi:10.1093/jnci/86.17.1352

10. Rauscher GH, Campbell RT, Wiley EL, Hoskins K, Stolley MR, Warnecke RB. Mediation of Racial and Ethnic Disparities in Estrogen/Progesterone Receptor-Negative Breast Cancer by Socioeconomic Position and Reproductive Factors. Am J Epidemiol. 2016;183(10):884–893. doi:10.1093/aje/kwv226

11. Warner ET, Tamimi RM, Hughes ME, et al. Racial and Ethnic Differences in Breast Cancer Survival: Mediating Effect of Tumor Characteristics and Sociodemographic and Treatment Factors. J Clin Oncol Off J Am Soc Clin Oncol. 2015;33(20):2254–2261. doi:10.1200/JCO.2014.57.1349

12. Costantino NS, Freeman B, Shriver CD, Ellsworth RE. Outcome Disparities in African American Compared with European American Women with ER+HER2-Tumors Treated within an Equal-Access Health Care System. Ethn Dis. 26(3):407–416. doi:10.18865/ed.26.3.407

13. Lovejoy LA, Eaglehouse YL, Hueman MT, Mostoller BJ, Shriver CD, Ellsworth RE. Evaluation of Surgical Disparities Between African American and European American Women Treated for Breast Cancer Within an Equal-Access Military Hospital. Ann Surg Oncol. 2019;26(12):3838–3845. doi:10.1245/s10434-019-07706-z

14. Watlington AT, Byers T, Mouchawar J, Sauaia A, Ellis J. Does having insurance affect differences in clinical presentation between Hispanic and non-Hispanic white women with breast cancer? Cancer. 2007;109(10):2093–2099. doi:10.1002/cncr.22640

15. Wojcik BE, Spinks MK, Stein CR. Effects of Screening Mammography on the Comparative Survival Rates of African American, White, and Hispanic Beneficiaries of a Comprehensive Health Care System. Breast J. 2003;9(3):175–183. doi:10.1046/j.1524-4741.2003.09308.x

16. Bao P-P, Cai H, Peng P, et al. Body mass index and weight change in relation to triple-negative breast cancer survival. Cancer Causes Control CCC. 2016;27(2):229–236. doi:10.1007/s10552-015-0700-7

17. Torio CM, Klassen AC, Curriero FC, Caballero B, Helzlsouer K. The Modifying Effect of Social Class on the Relationship Between Body Mass Index and Breast Cancer Incidence. Am J Public Health. 2010;100(1):146–151. doi:10.2105/AJPH.2007.126979

18. Troester MA, Sun X, Allott EH, et al. Racial Differences in PAM50 Subtypes in the Carolina Breast Cancer Study. J Natl Cancer Inst. 2018;110(2). doi:10.1093/jnci/djx135

19. Comparison of Breast Cancer Molecular Features and Survival by African and European Ancestry in The Cancer Genome Atlas | Breast Cancer | JAMA Oncology | JAMA Network. Accessed October 20, 2020. https://jamanetwork.com/journals/jamaoncology/fullarticle/2624532

20. Sparano JA, Wang M, Zhao F, et al. Race and Hormone Receptor–Positive Breast Cancer Outcomes in a Randomized Chemotherapy Trial. JNCI J Natl Cancer Inst. 2012;104(5):406–414. doi:10.1093/jnci/djr543

21. Haricharan S, Bainbridge MN, Scheet P, Brown PH. Somatic mutation load of estrogen receptor-positive breast tumors predicts overall survival: an analysis of genome sequence data. Breast Cancer Res Treat. 2014;146(1):211–220. doi:10.1007/s10549-014-2991-x

22. Haricharan S, Punturi N, Singh P, et al. Loss of MutL Disrupts Chk2-dependent Cell Cycle Control Through CDK4/6 to Promote Intrinsic Endocrine Therapy Resistance in Primary Breast Cancer. Cancer Discov. Published online August 11, 2017. doi:10.1158/2159-8290.CD-16-1179

23. Anurag M, Punturi N, Hoog J, Bainbridge MN, Ellis MJ, Haricharan S. Comprehensive Profiling of DNA Repair Defects in Breast Cancer Identifies a Novel Class of Endocrine Therapy Resistance Drivers. Clin Cancer Res Off J Am Assoc Cancer Res. 2018;24(19):4887–4899. doi:10.1158/1078-0432.CCR-17-3702

24. Danforth DN. Disparities in breast cancer outcomes between Caucasian and African American women: a model for describing the relationship of biological and nonbiological factors. Breast Cancer Res BCR. 2013;15(3):208. doi:10.1186/bcr3429

25. D’Arcy M, Fleming J, Robinson WR, Kirk EL, Perou CM, Troester MA. Race-associated biological differences among Luminal A breast tumors. Breast Cancer Res Treat. 2015;152(2):437–448. doi:10.1007/s10549-015-3474-4

26. cBioPortal for Cancer Genomics. Accessed October 26, 2020. https://www.cbioportal.org/study/summary?id=brca_tcga

27. Cerami E, Gao J, Dogrusoz U, et al. The cBio Cancer Genomics Portal: An Open Platform for Exploring Multidimensional Cancer Genomics Data. Cancer Discov. 2012;2(5):401–404. doi:10.1158/2159-8290.CD-12-0095

28. Pirone JR, D’Arcy M, Stewart DA, et al. Age-Associated Gene Expression in Normal Breast Tissue Mirrors Qualitative Age-at-Incidence Patterns for Breast Cancer. Cancer Epidemiol Prev Biomark. 2012;21(10):1735–1744. doi:10.1158/1055-9965.EPI-12-0451

29. Casbas-Hernandez P, Sun X, Roman-Perez E, et al. Tumor Intrinsic Subtype Is Reflected in Cancer-Adjacent Tissue. Cancer Epidemiol Prev Biomark. 2015;24(2):406–414. doi:10.1158/1055-9965.EPI-14-0934

30. Anurag M, Haricharan S, Ellis MJ. CDK4/6 Inhibitor Biomarker Research: Are We Barking Up the Wrong Tree? Clin Cancer Res. 2020;26(1):3–5. doi:10.1158/1078-0432.CCR-19-3119

31. Jackson SP, Bartek J. The DNA-damage response in human biology and disease. Nature. 2009;461(7267):1071–1078. doi:10.1038/nature08467

32. Dasika GK, Lin S-CJ, Zhao S, Sung P, Tomkinson A, Lee EY-HP. DNA damage-induced cell cycle checkpoints and DNA strand break repair in development and tumorigenesis. Oncogene. 1999;18(55):7883–7899. doi:10.1038/sj.onc.1203283

33. Shen D, Wu Y, Subbarao M, Bhat H, Chillar R, Vadgama JV. Mutation analysis of BRCA1 gene in African-American patients with breast cancer. J Natl Med Assoc. 2000;92(1):29–35.

34. Smith ZL, Eggener SE, Murphy AB. African-American Prostate Cancer Disparities. Curr Urol Rep. 2017;18(10):81. doi:10.1007/s11934-017-0724-5

35. Peres LC, Schildkraut JM. Racial/ethnic disparities in ovarian cancer research. Adv Cancer Res. 2020;146:1–21. doi:10.1016/bs.acr.2020.01.002

36. Adams SA, Fleming A, Brandt HM, et al. Racial disparities in cervical cancer mortality in an African American and European American cohort in South Carolina. J S C Med Assoc 1975. 2009;105(7):237–244.

37. Hollenbeck BK, Dunn RL, Ye Z, Hollingsworth JM, Lee CT, Birkmeyer JD. Racial differences in treatment and outcomes among patients with early stage bladder cancer. Cancer. 2010;116(1):50–56. doi:10.1002/cncr.24701

38. Brookfield KF, Cheung MC, Gomez C, et al. Survival disparities among African American women with invasive bladder cancer in Florida. Cancer. 2009;115(18):4196–4209. doi:10.1002/cncr.24497

