## Supplementary Mazumder_et_al_2020 for "Race specific differences in DNA damage repair dysregulation in breast cancer and association with outcome"

Supplementary Material:

Figure S1

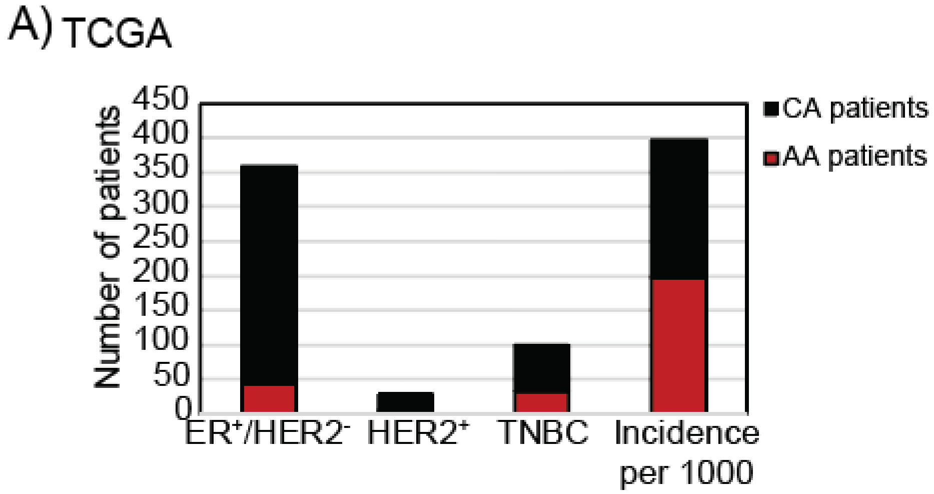


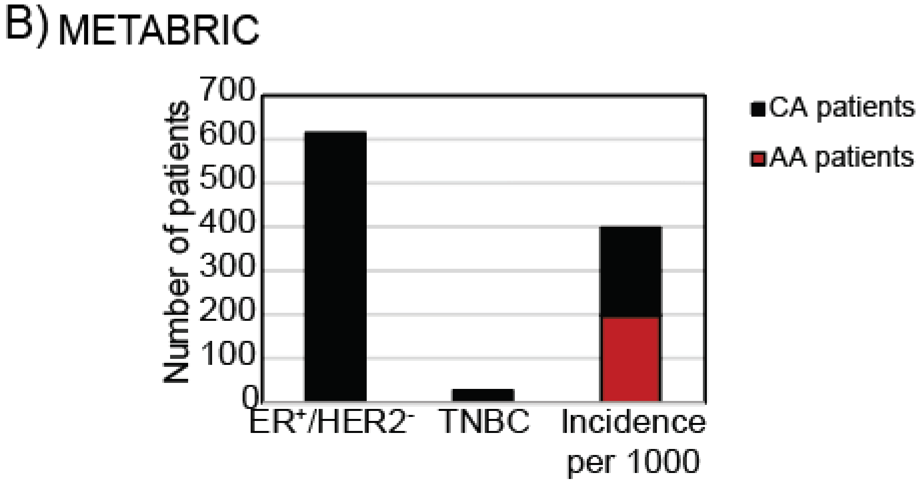

Figure S2

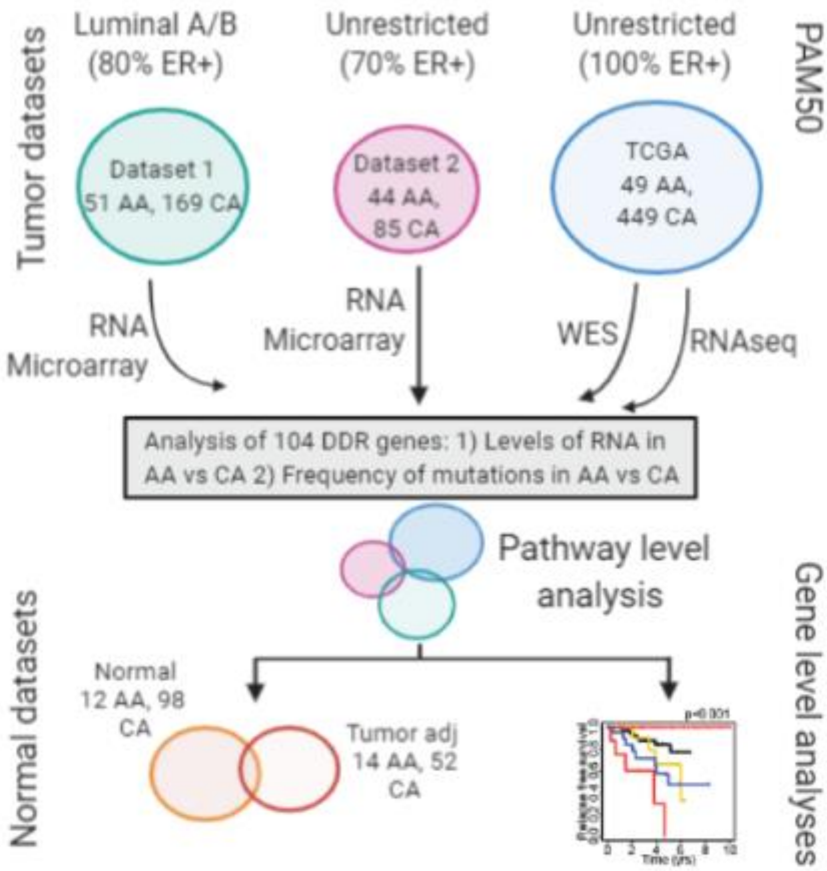

Tumor datasets
Luminal A/B
(80% ER+)
Unrestricted
(70% ER+)
Unrestricted
(100% ER+)
PAM50
Dataset 1
51 AA, 169 CA
Dataset 2
44 AA,
85 CA
TCGA
49 AA,
449 CA
RNA
Microarray
RNA
Microarray
WES
RNAseq
Analysis of 104 DDR genes: 1) Levels of RNA in AA vs CA 2) Frequency of mutations in AA vs CA
Pathway level
analysis
Normal datasets
Normal
12 AA, 98
CA
Tumor adj
14 AA, 52
CA
Gene level analyses

Figure S3

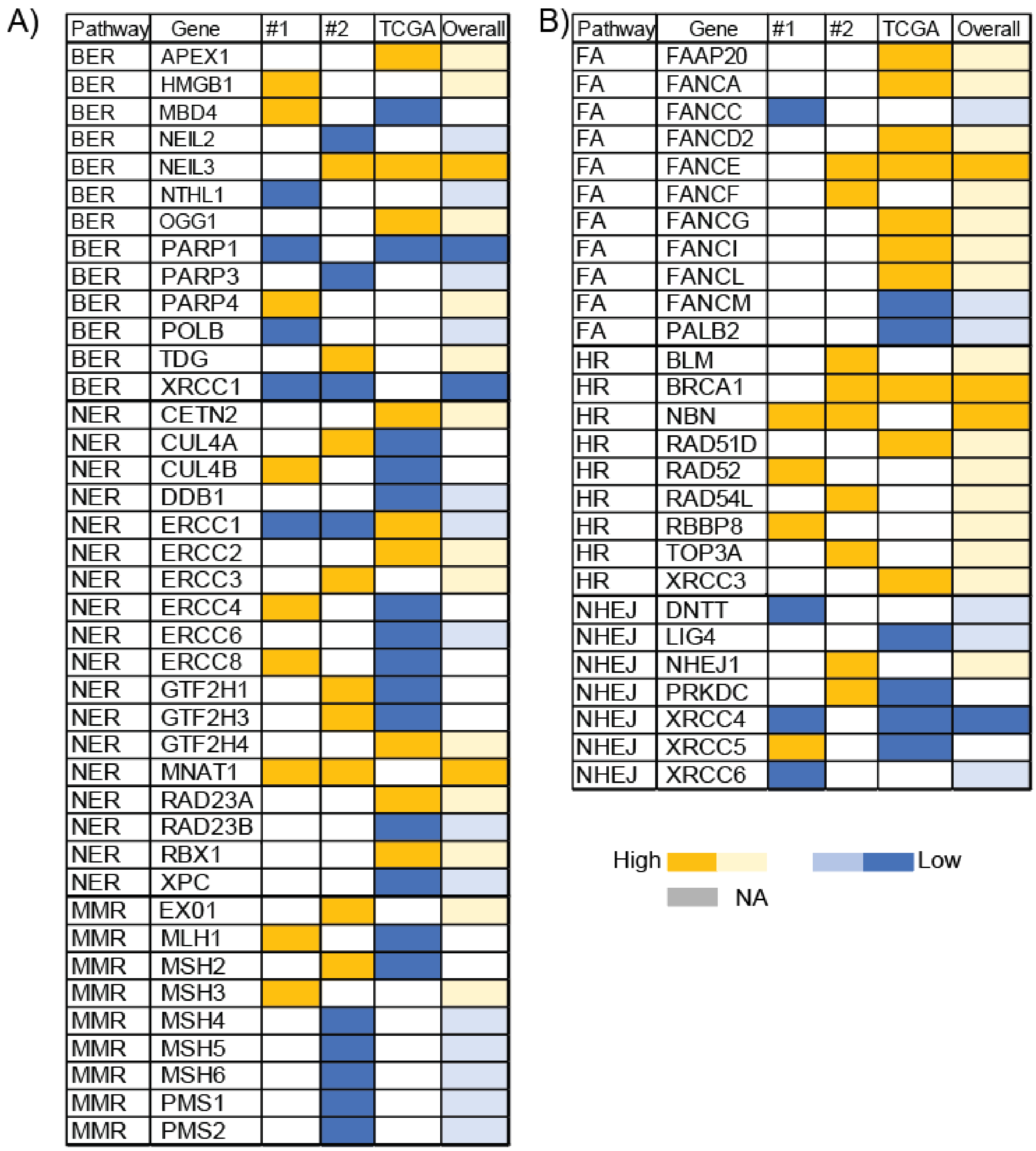
A)
Pathway
Gene
#1
#2
TCGA
Overall
BER APEX1
BER HMGB1
BER MBD4
BER NEIL2
BER NEIL3
BER NTHL1
BER OGG1
BER PARP1
BER PARP3
BER PARP4
BER POLB
BER TDG
BER XRCC1
NER CETN2
NER CUL4A
NER CUL4B
NER DDB1
NER ERCC1
NER ERCC2
NER ERCC3
NER ERCC4
NER ERCC6
NER ERCC8
NER GTF2H1
NER GTF2H3
NER GTF2H4
NER MNAT1
NER RAD23A
NER RAD23B
NER RBX1
NER XPC
MMR EX01
MMR MLH1
MMR MSH2
MMR MSH3
MMR MSH4
MMR MSH5
MMR MSH6
MMR PMS1
MMR PMS2
B)
Pathway
Gene
#1
#2
TCGA
Overall
FA FAAP20
FA FANCA
FA FANCC
FA FANCD2
FA FANCE
FA FANCF
FA FANCG
FA FANCI
FA FANCL
FA FANCM
FA PALB2
HR BLM
HR BRCA1
HR NBN
HR RAD51D
HR RAD52
HR RAD54L
HR RBBP8
HR TOP3A
HR XRCC3
NHEJ DNTT
NHEJ LIG4
NHEJ NHEJ1
NHEJ PRKDC
NHEJ XRCC4
NHEJ XRCC5
NHEJ XRCC6
High
Low
NA

Figure S4

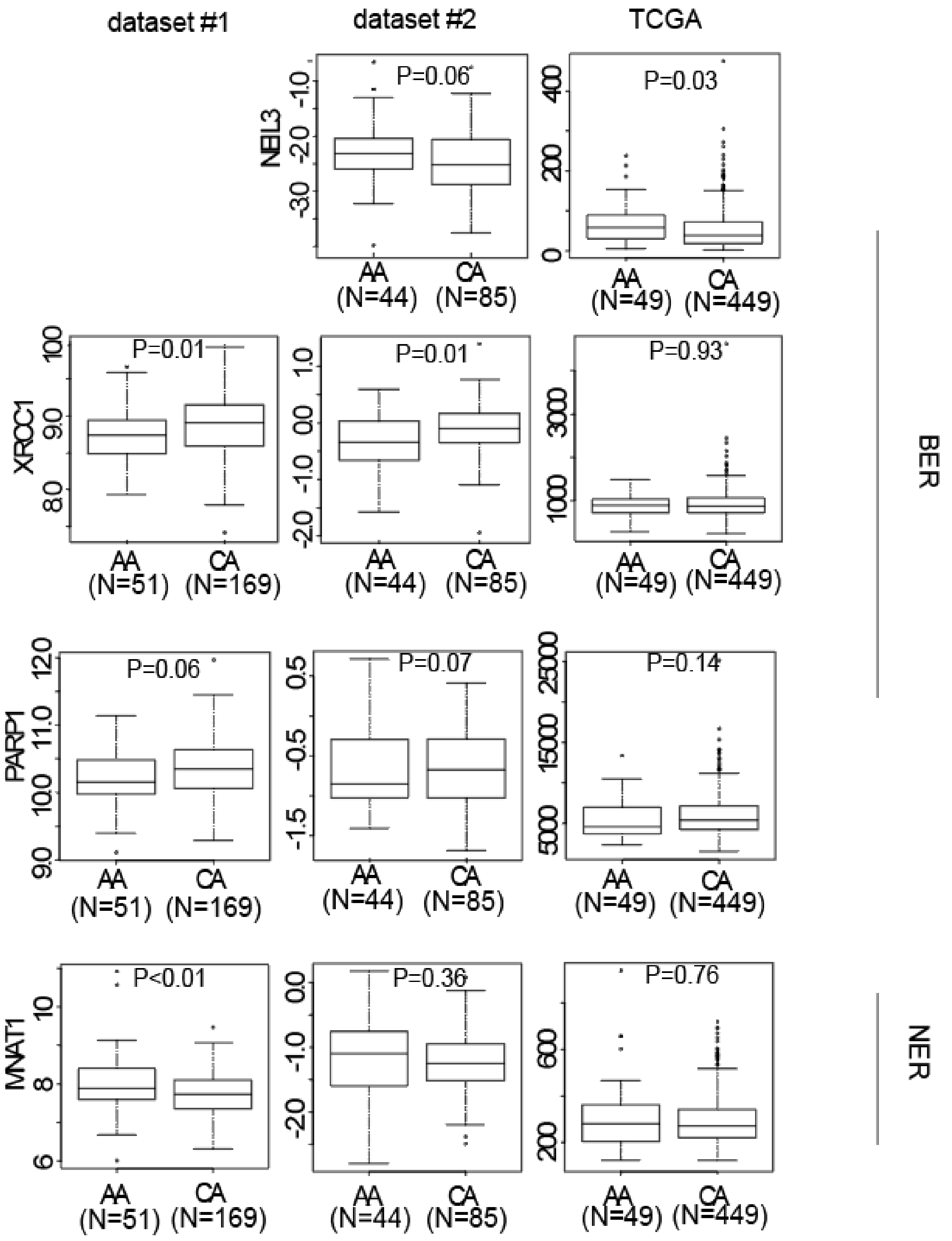

dataset #1
dataset #2
TCGA
NEIL3
P=0.06
P=0.03
XRCC1
P=0.01
P=0.01
P=0.93
PARP1
P=0.06
P=0.07
P=0.14
MNAT1
P<0.01
P=0.36
P=0.76
AA (N=51)
CA (N=169)
AA (N=44)
CA (N=85)
AA (N=49)
CA (N=449)
BER
NER

Figure S5

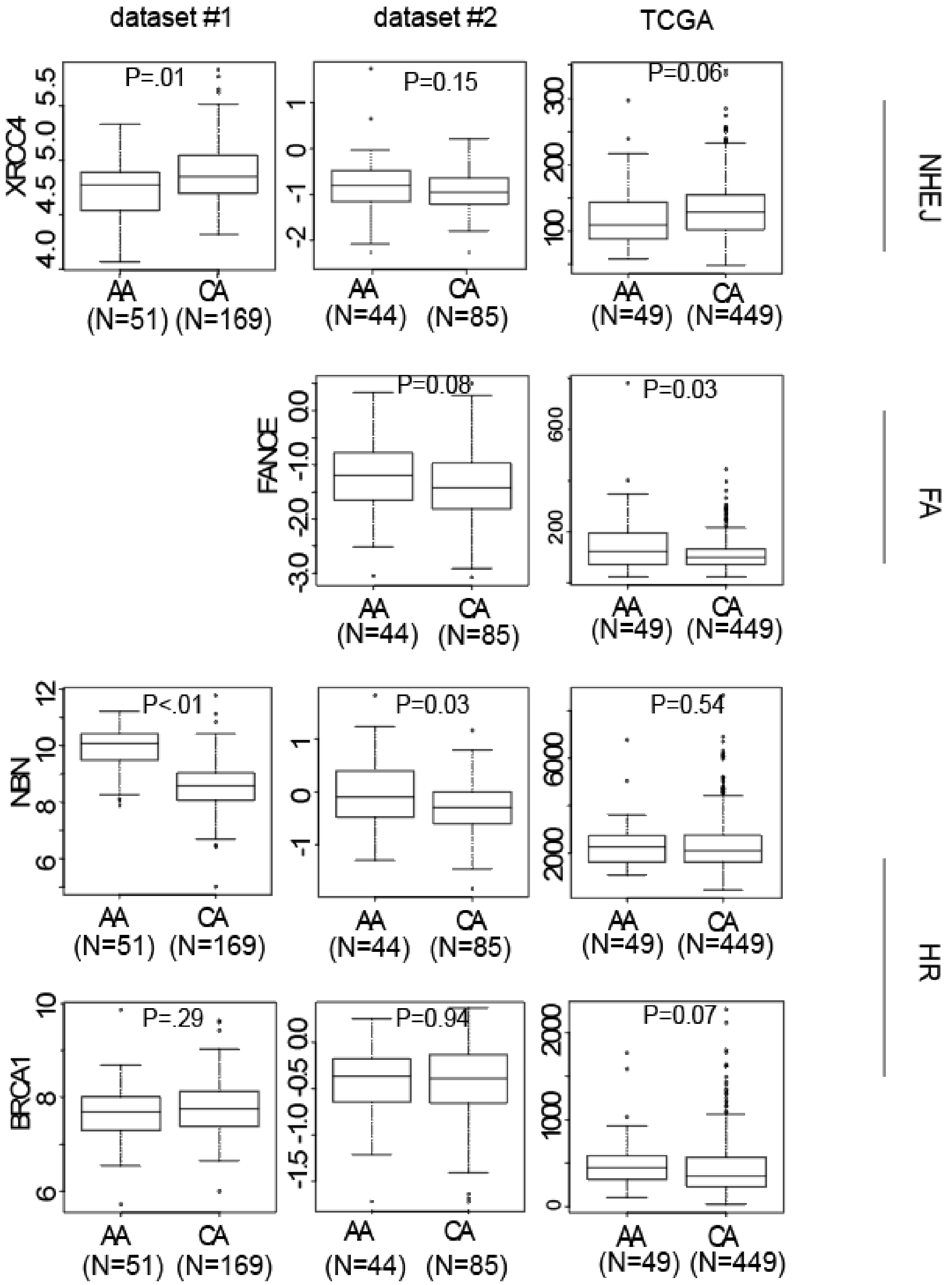
dataset #1
dataset #2
TCGA
XRCC4
FANCE
NBN
BRCA1
P=.01
P=0.15
P=0.06
P=0.08
P=0.03
P<.01
P=0.03
P=0.54
P=.29
P=0.94
P=0.07
AA (N=51)
CA (N=169)
AA (N=44)
CA (N=85)
AA (N=49)
CA (N=449)
NHEJ
FA
HR

Figure S6:

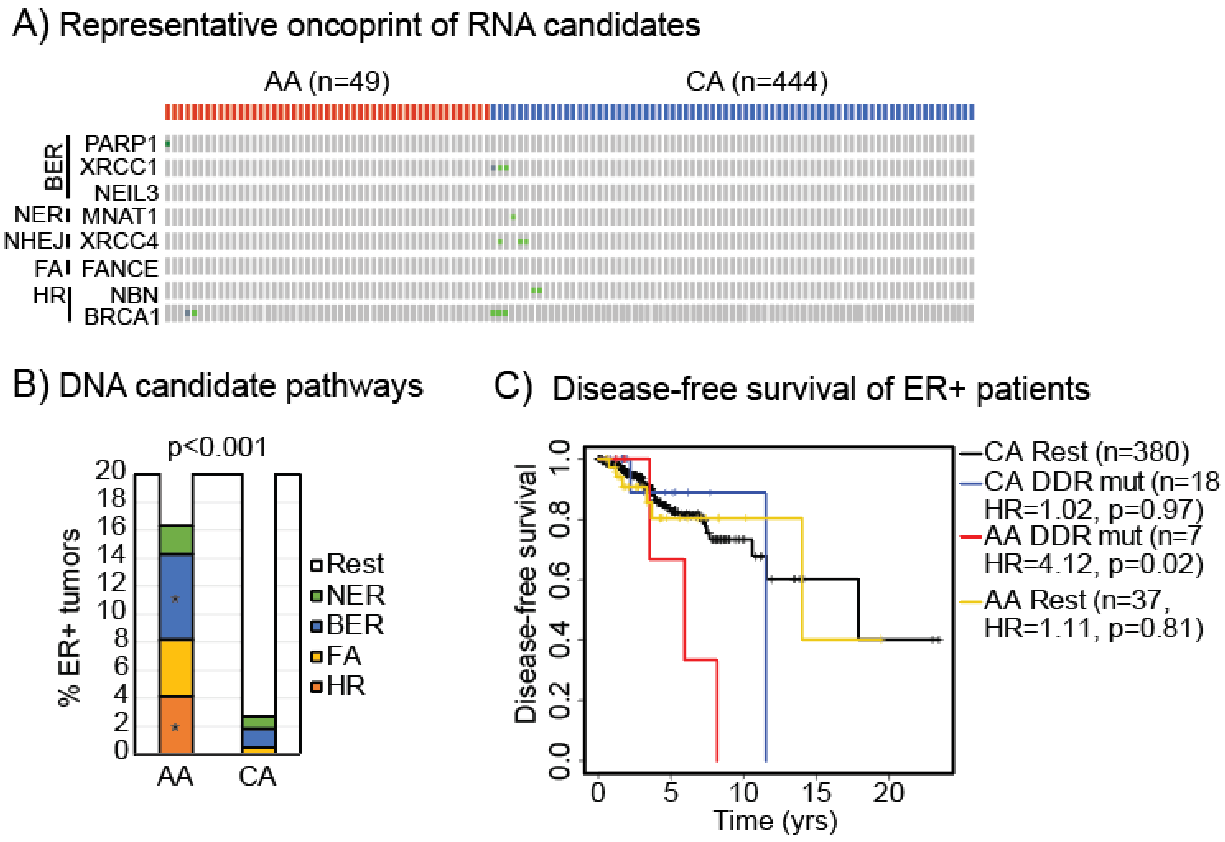

A) Representative oncoprint of RNA candidates
AA (n=49)
CA (n=444)
BER
PARP1
XRCC1
NEIL3
NER
MNAT1
NHEJ
XRCC4
FA
FANCE
HR
NBN
BRCA1
B) DNA candidate pathways
p<0.001
% ER+ tumors
20
18
16
14
12
10
8
6
4
2
0
AA
CA
Rest
NER
BER
FA
HR
C) Disease-free survival of ER+ patients
Disease-free survival
0.0
0.2
0.4
0.6
0.8
1.0
0
5
10
15
20
Time (yrs)
CA Rest (n=380)
CA DDR mut (n=18
HR=1.02, p=0.97)
AA DDR mut (n=7
HR=4.12, p=0.02)
AA Rest (n=37,
HR=1.11, p=0.81)

Figure S7

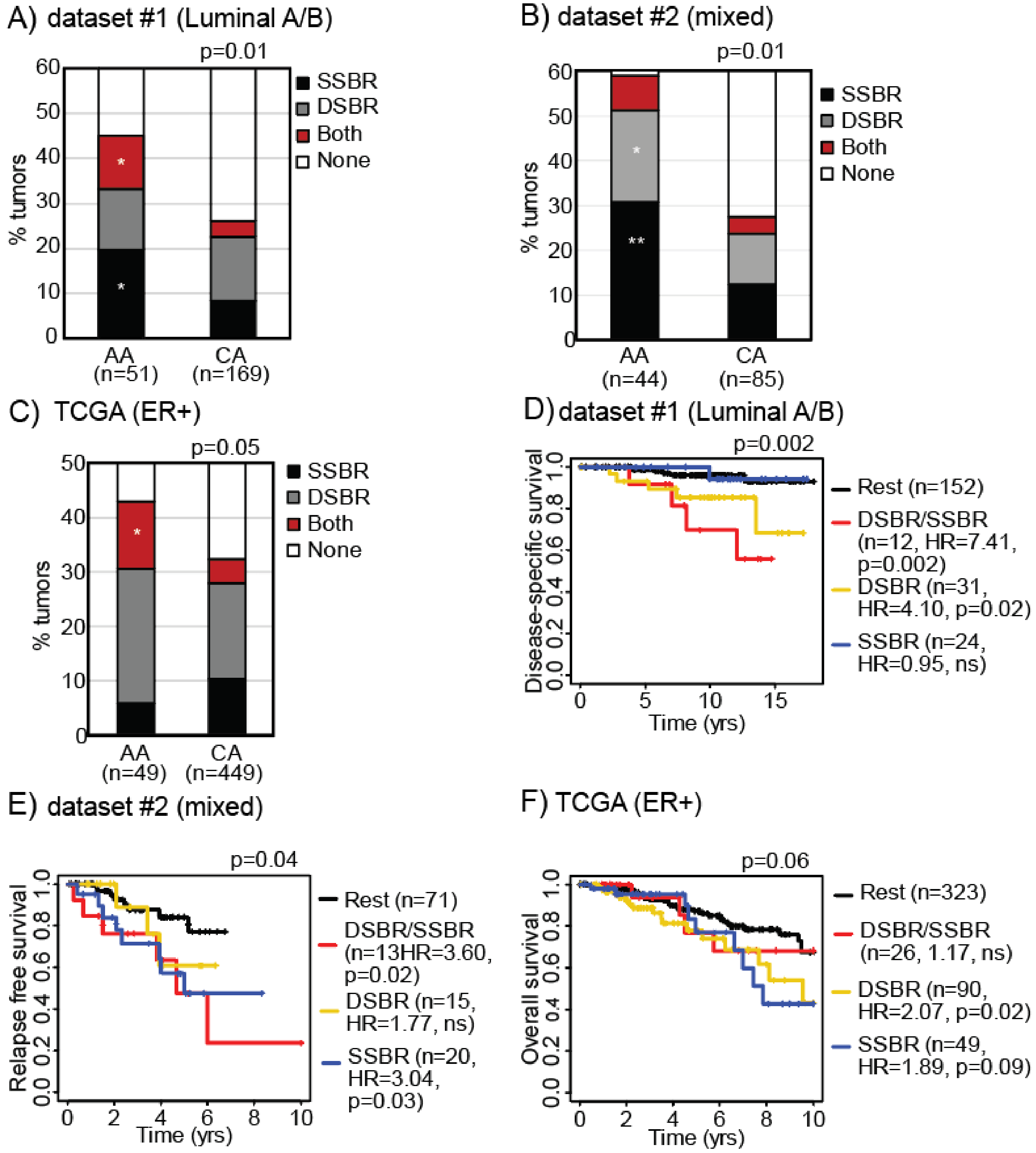

A) dataset #1 (Luminal A/B)
p=0.01
% tumors
AA (n=51)
CA (n=169)
SSBR
DSBR
Both
None
B) dataset #2 (mixed)
p=0.01
% tumors
AA (n=44)
CA (n=85)
SSBR
DSBR
Both
None
C) TCGA (ER+)
p=0.05
% tumors
AA (n=49)
CA (n=449)
SSBR
DSBR
Both
None
D) dataset #1 (Luminal A/B)
p=0.002
Disease-specific survival
Time (yrs)
Rest (n=152)
DSBR/SSBR (n=12, HR=7.41, p=0.002)
DSBR (n=31, HR=4.10, p=0.02)
SSBR (n=24, HR=0.95, ns)
E) dataset #2 (mixed)
p=0.04
Relapse free survival
Time (yrs)
Rest (n=71)
DSBR/SSBR (n=13HR=3.60, p=0.02)
DSBR (n=15, HR=1.77, ns)
SSBR (n=20, HR=3.04, p=0.03)
F) TCGA (ER+)
p=0.06
Overall survival
Time (yrs)
Rest (n=323)
DSBR/SSBR (n=26, 1.17, ns)
DSBR (n=90, HR=2.07, p=0.02)
SSBR (n=49, HR=1.89, p=0.09)

Table S1. List of genes with Functional Categories.

| Gene Name | DNA repair Pathway | Function |
|---|---|---|
| XPC | Nucleotide excision repair | Lesion recognition, recruits ATM/ATR |
| XPA | | Acts as scaffold for ATR to bind |
| RBX1 | | Lesion recognition |
| RAD23B | | Lesion recognition |
| RAD23A | | Lesion recognition |
| MNAT1 | | Acts as scaffold associates with ATM |
| GTF2H5 | | Acts as scaffold associates with ATM |
| GTF2H4 | | Acts as scaffold associates with ATM |
| GTF2H3 | | Acts as scaffold associates with ATM |
| GTF2H2 | | Acts as scaffold associates with ATM |
| GTF2H1 | | Acts as scaffold associates with ATM |
| ERCC8 | | Lesion recognition |
| ERCC6 | | Lesion recognition |
| ERCC5 | | Endonuclease/Repair activity |
| ERCC4 | | Endonuclease/Repair activity |
| ERCC3 | | Acts as Scaffold |
| ERCC2 | | Acts as scaffold |
| ERCC1 | | Endonuclease/Repair activity |
| DDB2 | | Lesion recognition/recruits ATM/ATR |
| DDB1 | | Lesion recognition/recruits ATM/ATR |
| CUL4B | | Lesion recognition/recruits ATM/ATR |
| CUL4A | | Lesion recognition/recruits ATM/ATR |
| CETN2 | | Lesion recognition/recruits ATM/ATR |
| XRCC1 | Base Excision Repair | Associates with CHK2 |

Table S1. List of genes with Functional Categories

| Gene Name | DNA repair Pathway | Function |
|---|---|---|
| UNG | | Lesion recognition |
| TDG | | Lesion recognition |
| SMUG1 | | Lesion recognition/associates with ATM/CHK2 |
| POLB | | Repair activity |
| PNKP | | Repair activity |
| PARP4 | | Repair activity |
| PARP3 | | Repair activity |
| PARP2 | | Repair activity |
| PARP1 | | Lesion recognition/repair activity |
| OGG1 | | Lesion recognition |
| NTHL1 | | Lesion recognition |
| NEIL3 | | Lesion recognition |
| NEIL2 | | Lesion recognition |
| NEIL1 | | Lesion recognition |
| MUTYH | | Lesion recognition/associate with ATM/CHK2 |
| MPG | | Lesion recognition/associate with ATM/CHK2 |
| MBD4 | | Lesion recognition |
| LIG3 | | Repair activity |
| HMGB1P4 | | Downstream effectors |
| HMGB1P1 | | Downstream effectors |
| HMGB1 | | Downstream effectors |
| APLF | | Downstream effectors |
| APEX2 | | Repair activity |
| APEX1 | | Repair activity |
| PMS2P3 | Mismatch repair pathway | Scaffold |
| PMS2 | | Scaffold/Repair activity |
| PMS1 | | Scaffold |
| MSH6 | | Lesion recognition/Scaffold associates with CHK2 |
| MSH5 | | Lesion Recognition |
| MSH4 | | Lesion Recognition |
| MSH3 | | Lesion Recognition/Scaffold associates with CHK2 |
| MSH2 | | Lesion Recognition/scaffold associates with CHK2 |
| MLH3 | | Lesion Recognition |

Table S1. List of genes with Functional Categories.

| Gene Name | DNA repair Pathway | Function |
|---|---|---|
| MLH1 | | Repair activity/Scaffold associates with ATM |
| EXO1 | | Scaffold |
| XRCC6 | Non-Homologous End Joining | Lesion recognition |
| XRCC5 | | Lesion recognition |
| XRCC4 | | Scaffold |
| PRKDC | | Scaffold |
| POLM | | Downstream effectors |
| NHEJ1 | | Scaffold |
| LIG4 | | Scaffold/associates with ATM |
| DNTT | | Downstream effectors |
| XRCC3 | Homologous recombination | Scaffold/Repair activity |
| XRCC2 | | Scaffold |
| TOP3B | | Downstream effectors |
| TOP3A | | Downstream effectors |
| SHFM1 | | Scaffold |
| RBBP8 | | Repair activity/endonuclease |
| RAD54L | | Lesion recognition |
| RAD54B | | Lesion recognition |
| RAD52 | | Repair activity |
| RAD51D | | Scaffold |
| RAD51B | | Scaffold |
| RAD51 | | Scaffold |
| NBN | | Lesion recognition/associates with ATM |
| MUS81 | | Repair activity/endonuclease |
| SLX1B | | Repair activity/endonuclease |
| SLX1A | | Repair activity/endonuclease |
| GEN1 | | Repair activity/endonuclease |
| EME2 | | Scaffold/Repair activity |
| EME1 | | Repair activity/endonuclease |
| DMC1 | | Repair activity/recombinases |
| BRCA1 | | Lesion recognition/Scaffold |
| BLM | | Repair activity |
| SLX4 | Fanconi Anemia | Scaffold |
| PALB2 | | Downstream effectors |

Table S1. List of genes with Functional Categories.

| Gene Name | DNA repair Pathway | Function |
|---|---|---|
| FANCM | | Scaffold/associates with ATR/CHK1 signaling |
| FANCL | | Scaffold/associates with ATR/CHK1 signaling |
| FANCI | | Scaffold/associates with ATR/CHK1 signaling |
| FANCG | | Scaffold/associates with ATR/CHK1 signaling |
| FANCF | | Scaffold/associates with ATR/CHK1 signaling |
| FANCE | | Scaffold/associates with ATR/CHK1 signaling |
| FANCD2 | | Repair activity |
| FANCC | | Scaffold/associates with ATR/CHK1 signaling |
| FANCB | | Scaffold |
| FANCA | | Scaffold/associates with ATR/CHK1 signaling |
| FAAP24 | | Lesion recognition |
| FAAP20 | | Scaffold/associates with ATR/CHK1 signaling |
| BRIP1 | | Scaffold |

Table S2. Patient characteristics from dataset #1

| Characteristic | CA N=169(%) | AA N=51(%) | p value |
|---|---|---|---|
| **Age** | | | |
| <40 | 14 (8.3) | 5 (9.8) | |
| 40-49 | 40 (23.7) | 13 (25.5) | |
| >=50 | 114 (67.5) | 33 (64.7) | |
| Missing | 1 (.5) | 0 | 0.92 |
| **Tumor Stage** | | | |
| I | 75 (44.4) | 23 (45.1) | |
| II | 73 (43.2) | 22 (43.1) | |
| III | 15 (8.9) | 5 (9.8) | |
| IV | 5 (3) | 1 (2) | |
| Unknown | 1 (.5) | 0 | 0.97 |
| **Tumor Grade** | | | |
| Well | 57 (33.7) | 9 (17.6) | |
| Moderate | 81 (47.9) | 30 (58.8) | |
| Poor | 30 (17.8) | 11 (21.6) | |
| Missing | 1 (.6) | 1 (2) | 0.14 |
| **ER status** | | | |
| Positive | 169 (100) | 50 (98) | |
| Negative | 0 | 1(2) | 0.07 |
| **PR status** | | | |
| Positive | 137 (81.1) | 39 (76.5) | |
| Negative | 31 (18.3) | 12 (23.5) | |
| Missing | 1 (.6) | 0 | 0.62 |
| **Subtype** | | | |
| Lum A | 141 (83.4) | 42 (82.4) | |
| Lum B | 28 (16.6) | 9 (17.6) | 0.86 |

Table S3. Patient characteristics of dataset #2

| Characteristic | CA N=85(%) | AA N=44 (%) | p value |
|---|---|---|---|
| **Age** | | | |
| <40 | 9 (10.6) | 5 (11.4) | |
| 40-49 | 20 (23.5) | 11 (25) | |
| >=50 | 54 (63.5) | 26 (59.1) | |
| Missing | 2 (2.4) | 2 (4.5) | 0.90 |
| **Node Status** | | | |
| Positive | 32 (37.6) | 24 (54.6) | |
| Negative | 51 (60) | 18 (40.9) | |
| Unknown | 2( 2.4) | 2 (4.5) | 0.11 |
| **Size** | | | |
| 1 | 23 (27.1) | 9 (20.5) | |
| 2 | 41 (48.2) | 20 (45.5) | |
| +3 | 19 (22.4) | 12 (27.2) | |
| Missing | 2 (2.3) | 3 (6.8) | 0.50 |
| **ER status** | | | |
| Positive | 57 (67.1) | 23 (52.3) | |
| Negative | 25 (29.4) | 19 (43.2) | |
| Missing | 3 (3.5) | 2 (4.5) | 0.26 |
| PR status | | | |
| Positive | 41 (48.2) | 19 (43.2) | |
| Negative | 34 (68) | 21 (47.7) | |
| Missing | 10 (11.8) | 4 (9.1) | 0.69 |
| **Her2 Status** | | | |
| Positive | 21 (24.7) | 11 (25) | |
| Negative | 58 (68.2) | 30 (68.2) | |
| Missing | 6 (7.1) | 3 (6.8) | 1.00 |

Table S4. Characteristics of TCGA dataset.

| Characteristic | CA N=449(%) | AA N=49 (%) | P value |
|---|---|---|---|
| **Age** | | | |
| <40 | 27 (6) | 7 (14.3) | |
| 40-49 | 89 (19.8) | 11 (22.4) | |
| >=50 | 333 (74.2) | 30 (61.2) | |
| Missing | 0 | 1 (2) | .002 |
| **Node Status** | | | |
| 0 | 2 (.4) | 0 | |
| 1 | 44 (9.8) | 4 (8.2) | |
| 2 | 55 (12.2) | 5 (10.2) | |
| +3 | 341 (75.9) | 37 (75.5) | |
| Missing | 7 (1.6) | 3 (6.1) | 0.28 |
| **ER status** | | | |
| Positive | 449 (100) | 49 (100) | |
| **PR status** | | | |
| Positive | 384 (85.5) | 40 (81.6) | |
| Negative | 62 (13.8) | 9 (18.4) | |
| Missing | 3 (.7) | 0 | 0.59 |
| **HER2 Status** | | | |
| Positive | 61 (13.6) | 3 (6.1) | |
| Negative | 240 (53.5) | 19 (38.8) | |
| Equivocal | 83 (18.5) | 15 (30.6) | |
| Missing | 65 (14.5) | 12 (24.5) | 0.02 |

Table S5. Cox proportional hazards for dataset #1

| Factor | HR | CI | p | n |
|---|---|---|---|---|
| DDR status | | | | |
| *Rest* | -- | | | 152 |
| *DSBR* | 2.02 | 0.53-7.61 | 0.30 | 31 |
| *SSBR* | 0.82 | 0.08-8.06 | 0.86 | 24 |
| *HR/SSBR** | 6.44 | 1.32-31.31 | 0.02 | 9 |
| *Other/SSBR* | 1.82 | 0.16-20.88 | 0.63 | 3 |
| PAM50 | | | | |
| *LumA* | -- | | | 182 |
| *LumB* | 0.92 | 0.23-3.67 | 0.90 | 37 |
| Tumor stage | | | | |
| *i* | -- | | | 97 |
| *ii* | 1.42 | 0.30-6.63 | 0.66 | 95 |
| *iii+** | 11.9 | 2.77-50.73 | <0.001 | 26 |
| Age | 1.01 | 0.97-1.05 | 0.67 | 218 |
| Race | | | | |
| *AA* | -- | | | 51 |
| *CA* | 0.78 | 0.22-2.78 | 0.71 | 168 |

Table S6. Cox proportional hazards for dataset #2

| Factor | HR | CI | p | n |
|---|---|---|---|---|
| DDR status | | | | |
| Rest | -- | | | 71 |
| DSBR | 0.63 | 0.19-2.92 | 0.51 | 16 |
| SSBR* | 4.38 | 1.25-15.85 | 0.03 | 19 |
| HR/SSBR*** | 16.84 | 2.92-73.13 | <.001 | 8 |
| PAM50 | | | | |
| LumA | -- | | | 50 |
| LumB' | 5.21 | 1.25-33.21 | 0.05 | 28 |
| HER2*** | 22.41 | 3.47-133.69 | <.001 | 15 |
| Basal-like*** | 40.42 | 7.58-225.46 | <.001 | 17 |
| Claudin-low | 1.9 | 0.35-47.05 | 0.64 | 7 |
| Normal-like*** | 180.1 | 8.90-3839.05 | <.001 | 2 |
| Tumor size | | | | |
| 1 | -- | | | 32 |
| 2 | N/A | N/A | N/A | 57 |
| 3 | N/A | N/A | N/A | 28 |
| Unk | N/A | N/A | N/A | 2 |
| Node | 0.73 | 0.24-2.22 | 0.58 | 119 |
| Age | 1 | 0.96-1.04 | 0.88 | 119 |
| Race | | | | |
| AA | -- | | | 40 |
| CA | 0.92 | 0.33-2.54 | 0.87 | 79 |

Table S7. Cox proportional hazards for TCGA

| Factor | HR | CI | p | n |
|---|---|---|---|---|
| DDR status | | | | |
| *Rest* | -- | | | 328 |
| *DSBR** | 2.1 | 1.05-4.22 | 0.04 | 90 |
| *SSBR* | 1.68 | 0.05-3.25 | 0.22 | 49 |
| *HR/SSBR* | 1.77 | 0.22-14.12 | 0.59 | 9 |
| *Other/SSBR* | 0.41 | 0.05-3.25 | 0.40 | 17 |
| PR status | | | | |
| *Neg* | -- | | | 71 |
| *Pos* | 0.72 | 0.33-1.58 | 0.99 | 419 |
| HER2 status | | | | |
| *Neg* | -- | | | 345 |
| *Pos** | 2.09 | 1.11-3.95 | 0.02 | 110 |
| Tumor Stage | | | | |
| *1* | -- | | | 153 |
| 2 | 0.88 | 0.44-1.79 | 0.73 | 257 |
| *3* | 1.43 | 0.67-3.04 | 0.35 | 83 |
| Age | 1.02 | 0.99-1.04 | 0.15 | 492 |
| Race | | | | |
| *AA* | -- | | | 49 |
| *CA* | 1.91 | 0.45-8.16 | 0.38 | 444 |

Supplementary figure legends

**FigureS1: Major breast cancer patient tumor datasets underrepresent AAs.** Stacked column graphs representing the number of AA (red) and CA (black) patient tumors in two widely used breast cancer datasets, TCGA (A) and METABRIC (B). Numbers of AA and CA patients diagnosed with breast cancer per 1000 diagnoses in the USA in 2019 are included for comparison.

**Figure S2: Schematic of study.** Outline and details of datasets used in the study and order in which analyses were conducted and are described.

**FigureS3: Landscape of DDR dysregulation specific to AA tumors.** Heatmap showing all single stranded break repair (SSBR) (A) and double strand break repair (DSBR) (B) genes that had significantly different RNA levels in tumors from AA patients when compared to tumors from CAs in at least one dataset. Genes were considered to have significantly different RNA levels if they had a $q<0.25$ based on p-values generated by comparing RNA levels of 104 DDR genes unique to 6 common DDR pathways using Wilcoxon Rank Sum test. RNA levels are denoted as higher in AA tumors (yellow), lower (blue) or missing (grey). Supports data presented in **Figure 2**.

**FigureS4 and S5: Dysregulation of candidate DDR genes.** Boxplots showing RNA levels of each DDR gene that was dysregulated in at least two of the three tumor datasets analyzed. Wilcoxon Rank Sum test determined p-values. Horizontal line depicts the median and error bars the standard deviation. Supports data in **Figure 2**.

**FigureS6: Mutational landscape of DDR genes in AA tumors.** (A) Oncoprint from cBio depicting mutations in candidate genes identified in the analysis in **Figure 2** in AA (red) and CA (blue) ER+ patient tumors from TCGA. (B) Stacked column graph representing the proportion of ER+ tumors in AA vs CA patients with mutations in any DDR gene in the specified pathways. Pearson's Chi Squared test determined p-values. (C)Kaplan-Meier disease free survival curves for AA and CA ER+ patients whose tumors harbored mutations in any DDR gene. Analysis was restricted to genes that were mutated at least once in AA tumors. Log rank test determined p-values. HR, hazard ratios for survival curves, and homologous recombination in context of DDR pathways. Supports data in **Figure 2**.

**FigureS7: Association of AA-specific DDR dysregulation with patient outcomes.** (A-C) Stacked columns representing the percentage of AA vs CA tumors with dysregulation of any DDR genes from the specified pathways in dataset #1 (A), dataset #2 (B) and TCGA (C). Pearson's chi-squared test determined p-values for overall distribution of DDR gene dysregulation in AAs vs CAs, and also by individual pathways. Associated data presented in **Figure 4A-C**. (D-F) Kaplan-Meier survival curves representing disease-specific survival in dataset #1 (D), relapse-free survival in dataset #2 (E) and overall survival in TCGA (F) associated with patients whose tumors had dysregulation of specified candidate DDR genes by pathway relative to tumors that did not. Log rank test determined p-values. Associated data in **Figure 4D-F**, and Cox Regression Proportional Hazards assessment in **Tables S5-7**. HR in survival curves, hazard ratio, $p<0.05$, *; $p<0.01$, **; $p<0.001$, ***.
